# Organophosphorus diisopropylfluorophosphate (DFP) intoxication in zebrafish larvae causes behavioral defects, neuronal hyperexcitation and neuronal death

**DOI:** 10.1101/2019.12.15.876649

**Authors:** Alexandre Brenet, Julie Somkhit, Rahma Hassan-Abdi, Constantin Yanicostas, Christiane Romain, Olivier Bar, Alexandre Igert, Dominique Saurat, Nicolas Taudon, Gregory Dal-Bo, Florian Nachon, Nina Dupuis, Nadia Soussi-Yanicostas

## Abstract

With millions of intoxications each year and over 200,000 deaths, organophosphorus (OP) compounds are an important public health issue worldwide. OP poisoning induces cholinergic syndrome, with muscle weakness, hypertension, and neuron damage that may lead to epileptic seizures and permanent psychomotor deficits. Existing countermeasures are lifesaving but do not prevent long-lasting neuronal comorbidities, emphasizing the urgent need for animal models to better understand OP neurotoxicity and identify novel antidotes. Here, using diisopropylfluorophosphate (DFP), a prototypic and moderately toxic OP, combined with zebrafish larvae, we first showed that DFP poisoning caused major acetylcholinesterase inhibition, resulting in paralysis and CNS neuron hyperactivation, as indicated by increased neuronal calcium transients and overexpression of the immediate early genes *fosab, junBa, npas4b*, and *atf3*. In addition to these epileptiform seizure-like events, DFP-exposed larvae showed increased neuronal apoptosis, which were both partially alleviated by diazepam treatment, suggesting a causal link between neuronal hyperexcitation and cell death. Last, DFP poisoning induced an altered balance of glutamatergic/GABAergic synaptic activity with increased NR2B-NMDA receptor accumulation combined with decreased GAD65/67 and gephyrin protein accumulation. The zebrafish DFP model presented here thus provides important novel insights into the pathophysiology of OP intoxication, making it a promising model to identify novel antidotes.

## Introduction

Organophosphorus (OP) compounds comprise highly poisonous substances widely used as chemical pesticides but also as warfare agents. As a result of their massive use for agricultural purposes worldwide, OP poisoning represents a major public health issue with 3 million severe intoxications reported annually and more than 200,000 deaths, primarily suicides^1–4^. OPs are potent inhibitors of cholinesterases, including acetylcholinesterase (AChE), whose blockade causes a massive accumulation of acetylcholine and overstimulation of cholinergic receptors (ChRs) at both neuromuscular junctions and CNS cholinergic synapses^5^. In the brain, ChR hyperactivation causes epileptic seizures, which if not rapidly treated, may develop into life-threatening *status epilepticus* (SE)^6^. Besides their immediate toxicity, OPs also cause long-term neurological comorbidities, such as psychomotor deficits and recurrent seizures^7,8^. Existing countermeasures combine ChR agonist atropine with an AChE reactivator such as 2-PAM, often associated with □-aminobutyric acid receptor (GABAR) agonist diazepam (DZP). These treatments are lifesaving, but do not reverse brain damage or prevent subsequent occurrence of seizures or psychomotor deficits, emphasizing the need for new, fully efficient antidotes. To meet this need, an animal model of OP poisoning that would faithfully reproduce the consequences of OP intoxication in humans and be amenable to drug screening would help achieve a better understanding of the pathophysiology of OP poisoning and identify therapeutic entities counteracting the harmful effects of these compounds. Although it has long been known that acute OP intoxication causes neuropathological changes in the brain^8–11^, the precise nature of these changes and their extent remain under-researched.

Here, we describe a zebrafish model of OP poisoning and characterize the neuron defects induced by acute intoxication. Over the past decade, besides its rapidly expanding use as a human disease model^12–16^, the zebrafish has become one of the leading animal models for toxicology research and drug discovery^17^. The zebrafish is a versatile, powerful and easy-to-breed vertebrate model that offers significant advantages for *in vivo* drug screening and neurotoxicology investigations. These advantages include a CNS that displays an overall organization similar to that of mammals with full conservation of the different neuronal and glial cell types and all neurotransmitters^18–22^. To model OP poisoning in zebrafish, we chose DFP, an OP analogue of the warfare agent sarin, but less volatile and much less dangerous, that has been widely used as a prototypic OP in toxicology research. In particular, previous studies in rodents have shown that acute DFP poisoning causes potent AChE inhibition, inducing epileptic seizures, neuronal death, memory impairment and neuroinflammation^9,23^.

To better understand the physiopathology of OP poisoning and subsequent neuronal deficits, we combined behavioral analysis and *in vivo* recording of neuronal calcium transients with molecular and immunocytochemical approaches. Our data showed that zebrafish larvae exposed to DFP displayed major inhibition of AChE activity correlated with reduced motor activity. DFP poisoning also induced CNS hyperexcitation as shown by both the large increase in the number of calcium transients in brain neurons and the overexpression of the IEGs *fosab, junBa, npas4b*, and *atf3*. In addition to epileptiform seizures, DFP exposure also caused an increased neuronal apoptosis, both partially mitigated by diazepam administration, suggesting that the increased neuronal apoptosis was a direct consequence of neuronal hyperexcitation. Last, DFP poisoning caused increased NR2B-NMDA subunit receptor accumulation combined with decreased accumulation of both GAD65/67 and gephyrin proteins, suggesting that DFP exposure promotes epileptiform neuronal hyperexcitation as a result of a shift in the glutamatergic/GABAergic balance activity toward glutamate-mediated excitation.

The zebrafish model of DFP poisoning faithfully reproduces the neuronal deficits observed in both humans and rodents exposed to DFP, and also provides interesting new insights into the neurotoxicity of OP agents, making it a promising tool to identify novel, fully efficient antidotes.

## Results

### Larvae exposed to DFP showed increased mortality, reduced motility and AChE inhibition

As a prerequisite for establishing a zebrafish model of DFP poisoning, we first determined DFP toxicity in zebrafish *in vivo*, 5 days post-fertilization (dpf) larvae were exposed to 15, 20, 30, and 50 µM DFP and studied over a 24 h period. Results showed that all larvae exposed to 20 µM DFP or higher concentrations either died during the first 6 h of incubation or displayed major phenotypic defects, including curled tails and major reduction of head and eye volumes (Fig. **1b** and Supplementary Fig. **1)**. As our purpose was to investigate acute DFP neurotoxicity and subsequent brain damage, we selected 15 μM DFP and a 6 h exposure time, an experimental setup (Fig. **1a**) that caused a larval lethality of 20% (LC 20). Next, to measure DFP stability of this compound once diluted in fish water, a 15 µM solution was prepared by diluting a stock solution with fish water. Residual DFP concentrations were then determined at different time points over a 24 h period. Results showed that DFP diluted in fish water was almost stable, with an average loss per hour of approximately 2% (Fig. **1c**). Then, we analyzed the phenotype of larvae exposed to 15 μM DFP for 6 h. We observed no visible phenotypic defects in surviving larvae (Fig. **1d-e** and **f-i**), except a slight reduction of body length (Fig. **1i**). Histopathological analysis also indicated that following a 6 h exposure to 15 µM DFP, surviving larvae did not show any visible histological abnormalities in the brain (Supplementary Fig. **2**).

**Fig. 1.**
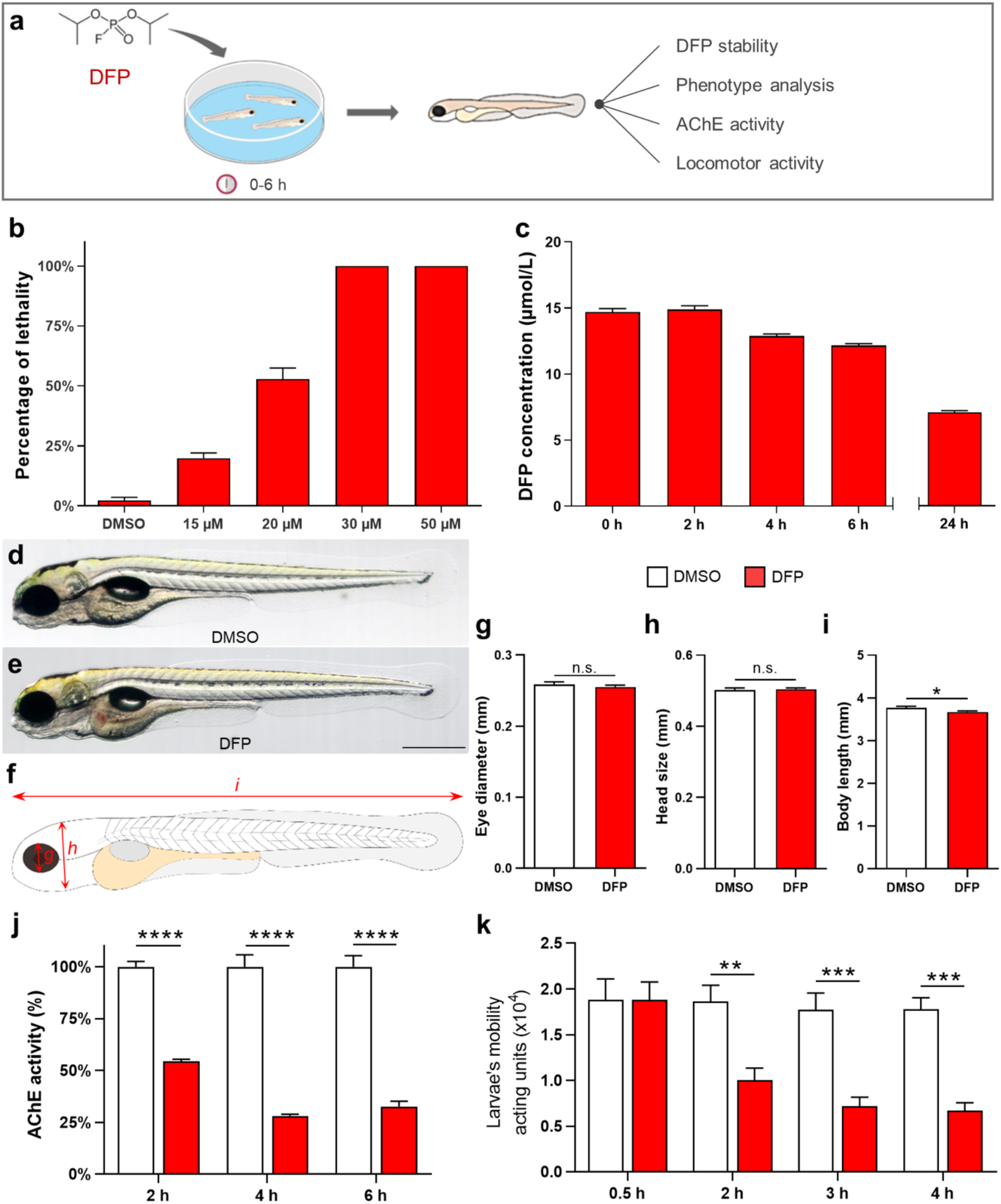
DFP-exposed zebrafish larvae displayed reduced motility and AChE inhibition. (**a**) As the experimental set-up, 5 dpf larvae were exposed to 15, 20, 30 or 50 µM DFP or vehicle (1% DMSO) and larval lethality, phenotypic defects, motor activity, and AChE activity were studied during the next 6 h. (**b**) Lethality rates of 5 dpf larvae exposed for 6 h to 15, 20, 30 or 50 µM DFP and selection of 15 µM DFP as optimal concentration (LC20). (**c**) Residual concentrations of DFP measured at different time points from a 15 µM solution diluted in fish water (E3 medium). (**d, e**) Morphology of 5 dpf larvae exposed for 6 h to either vehicle (**d**) or 15 µM DFP **(e**). (**f**) Scheme depicting measurements of zebrafish larva morphology. (**g-i**) Quantification of eye diameter (**g**), head size (**h**) and body length (**i**) in larvae exposed for 6 h to either vehicle (*n* = 26) or 15 µM DFP (*n* = 26) (Student’s unpaired *t*-test: n.s., non-significant; *, *p* < 0.05). (**j**) Quantification of AChE activity in larvae exposed to 15 µM DFP (*n* = 5) or vehicle (*n* = 5), for 2, 4, and 6 h (two-way ANOVA with Sidak’s multiple comparisons test: ****, *p* < 0.0001). (**k**) Motor activity of 5 dpf larvae exposed to either 15 µM DFP (*n* = 44) or vehicle (*n* = 12) (two-way ANOVA with Sidak’s multiple comparisons test: **, *p* < 0.01; ***, *p* < 0.001).

As AChE inhibition is the key hallmark of OP poisoning, we measured AChE activity in larvae exposed to 15 µM DFP, and we observed a 50% decrease in AChE activity as early as 2 h after DFP addition, worsening to 25% residual activity from 4 h exposure onwards (Fig. **1j**). We next investigated whether the decreased AChE activity was correlated with the paralysis of the larvae. We observed that DFP-treated larvae showed a marked decreased locomotor activity compared to control larvae (Fig. **1k**). Thus, after DFP exposure, zebrafish larvae displayed reduced motor activity as the likely result of a massive AChE inhibition.

### DFP exposure induced neuronal hyperexcitation

Increased transcription of the IEG *c-Fos* in brain neurons has been repeatedly observed following epileptic seizures.^24^ As a first investigation of the consequences of DFP poisoning on neuronal activity, we therefore studied, by immunocytochemistry, the accumulation of the Fosab protein, the zebrafish orthologue of c-Fos, in the brain of DFP-treated and control larvae (Fig. **2a, b**). Results clearly indicated that DFP-treated larvae displayed a highly significant increased number of neurons expressing Fosab in both the optic tectum (Fig. **2 c-e**), and telencephalon (Supplementary Fig. **3**). To further investigate whether DFP exposure induces overexpression of other IEGs, we analyzed, by qRT-PCR, the expression in the brain of *fosab, atf3, junBa, npas4a*, and *npas4b*, the zebrafish orthologues of *C-FOS, JUNB, ATF3*, and *NPAS4*, respectively, four genes that are expressed at high levels in the very first hours that follow epileptic seizures in pharmacological models of epilepsy in rats^25^. Interestingly, qRT-PCR showed a significant increase in the accumulation of *fosab* (control: 0.989 ± 0.088 *vs DFP*: 5.305 ± 0.804; *p* < 0.0001), *junBa* (control: 0.938 ± 0.197 *vs DFP*: 4.619 ± 0.603; *p* < 0.001), *atf3* (control: 0.993 ± 0.047 *vs DFP*: 5.723 ± 0.603; *p* < 0.003), and *npas4b* (control: 1.007 ± 0.054 *vs DFP*: 1.294 ± 0.109; *p* < 0.05), in larvae exposed to DFP when compared to that observed in controls (Fig. **2f**), suggesting that DFP exposure causes neuronal hyperexcitation. Note that expression of the second zebrafish orthologue of *NPAS4, npas4a*, did not change following DFP exposure (control: 1.03 ± 0.228 *vs DFP*: 1.226 ± 1.175; *p* = 0.5) (Fig. **2f**).

**Fig. 2.**
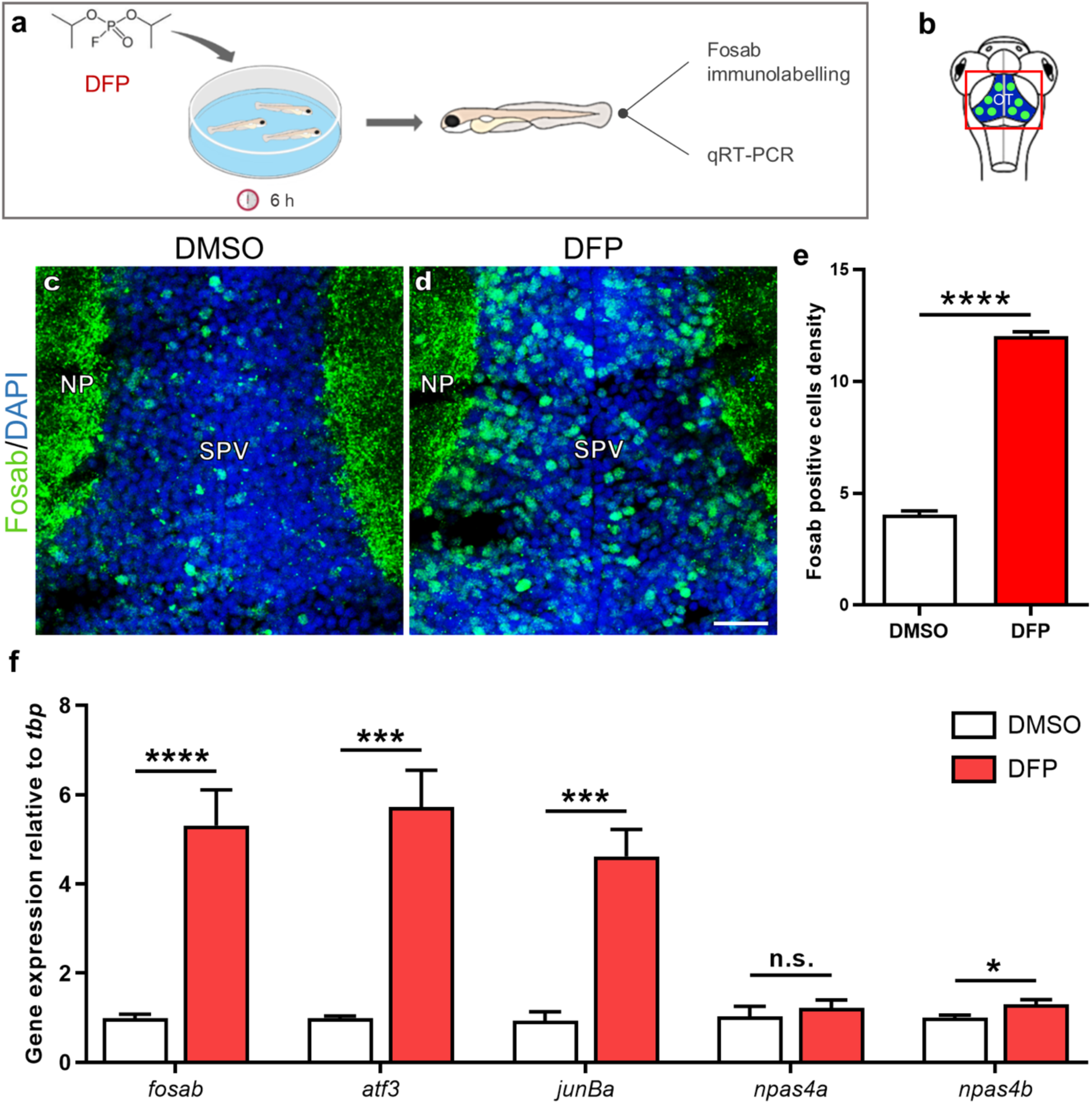
DFP exposure induces overexpression of the IEGs *fosab, atf3, junBa*, and *npas4b*. (**a**) As the experimental set-up, 5 dpf larvae were exposed for 6 h to either 15 µM DFP or vehicle (1% DMSO) before processing for either Fosab immunostaining or brain dissection followed by RNA extraction and qRT-PCR analysis. (**b**) Scheme of a 5 dpf larva’s head with the red box showing the region of interest in the brain, uncovering the optic tectum (OT). (**c**,**d**) Fosab immunolabeling of optic tectum neurons in 5 dpf larvae exposed to either vehicle (**c**) or 15 µM DFP (**d**). Scale bar: 20 µm. (**e**) Quantification of Fosab-expressing neuronal density in the optic tectum of 5 dpf larvae exposed to either vehicle (*N* = 3; *n* = 8) or 15 µM DFP (*N* = 3; *n* = 11) (unpaired *t*-test: ****, *p* < 0.0001). (**f**) qRT-PCR analysis of the accumulation of *fosab, atf3, junBa, npas4a* and *npas4b* RNAs relative to that of *tbp* in 5 dpf larvae exposed to either vehicle (*n* = 6) or 15 µM DFP (*n* = 6) (Student’s unpaired *t*-test: n.s., non-significant; *, *p* < 0.05; ***, *p* < 0.001; ****, *p* < 0.0001). *N* = number of larvae and *n* = number of slices. Abbreviations: NP, neuropil; SPV, stratum periventriculare.

We then sought to visualize the increased neuronal activity in larvae exposed to DFP using calcium imaging (Fig. **3a, b**), a technique that fully reflects neuronal activity in zebrafish epilepsy models *in vivo*^14,26^. Five dpf larvae from the transgenic line Tg[Huc:GCaMP5G] were treated with DFP or vehicle (DMSO) and calcium transients were recorded during the following 6 h using time-lapse confocal microscopy (Supplementary Videos **1** and **2**). Interestingly, in DFP-treated larvae, we observed numerous intense calcium transients, from approximately 1 h post-exposure onward, which were highly reminiscent of those seen in zebrafish genetic epilepsy models^14^. Moreover, these calcium transients were mostly seen in the neuropil region, where a large number of synaptic connections occur (Fig. **3c-f**). Quantification of calcium fluorescence signals confirmed that as early as 1 h after DFP addition, massive calcium transients were detected in DFP-treated larvae (Fig. **3g**). We also observed a significant increase in the frequency of the calcium transients from 2 h post-exposure onward (Fig. **3h**). Importantly, 3 h after DFP exposure, all larvae displayed numerous massive calcium transients, which were never observed in control larvae, strongly suggesting that zebrafish larvae exposed to acute DFP poisoning display massive neuronal hyperexcitation reminiscent of epileptic seizures. Diazepam (DZP), a GABA receptor agonist of the benzodiazepine family and a component of existing OP cocktail antidotes^27^, is a well-known inhibitor of neuronal excitation. We therefore checked whether administration of DZP was able to mitigate the increased number and intensity of calcium transients seen in DFP-exposed larvae. Data showed that massive calcium transients seen in larvae exposed for 5 h to 15 µM DFP were markedly reduced in the minutes that followed the addition of 40 µM DZP (Fig. **3i, j**), suggesting that the zebrafish DFP model faithfully reproduces the physiopathology of OP poisoning in humans.

**Fig. 3.**
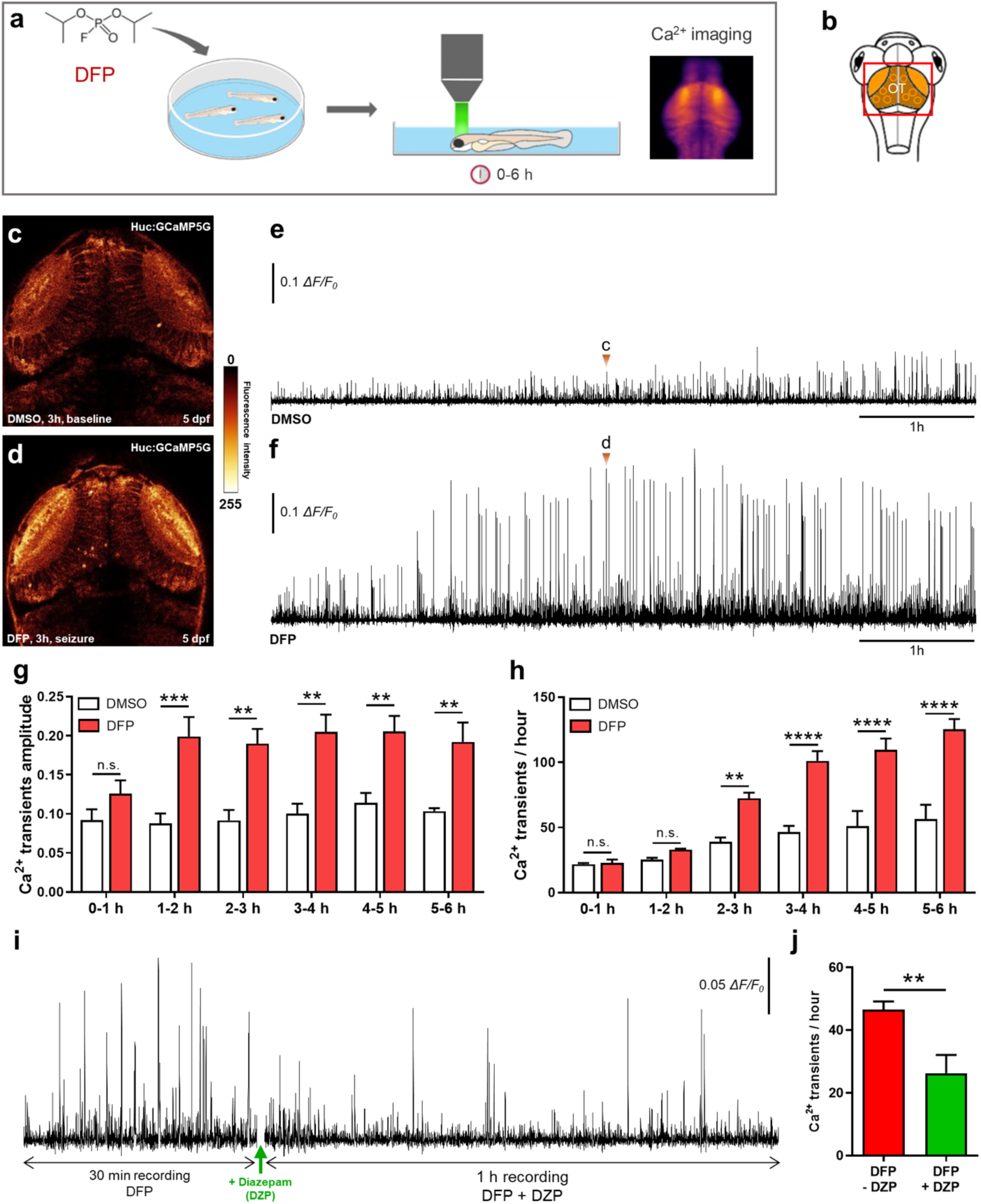
DFP exposure caused neuronal hyperexcitation. (**a**) As the experimental set-up, 5 dpf Tg[Huc:GCaMP5G] larvae were exposed to either 15 µM DFP or vehicle (1% DMSO), and calcium transients were recorded in brain neurons during the next 6 h. (**b**) Scheme of a 5 dpf larva’s head with the red box showing the region of interest in the brain, uncovering the optic tectum (OT). (**c-d**) Snapshot views of calcium imaging in a 5 dpf Tg[Huc:GCaMP5G] larva’s brain showing baseline calcium transients (**c** in Fig. **2e**) and seizure-like hyperactivity seen 3 h after exposure to 15 µM DFP (**d** in Fig. **2f**). (**e, f**) Baseline calcium transients detected in 5 dpf Tg[Huc:GCaMP5G] control larvae (*n* = 3) (**e**) and massive calcium transients detected in 5 dpf Tg[Huc:GCaMP5G] larvae exposed for 6 h to 15 µM DFP (*n* = 4) (**f**). (**g**) Amplitude of calcium transients detected in 5 dpf Tg[Huc:GCaMP5G] larvae at different time points following exposure to either 15 µM DFP (*n* = 4) or vehicle (*n* = 3) (two-way ANOVA with Sidak’s multiple comparisons test: n.s., non-significant; **, *p* < 0.01; ***, *p* < 0.001). (**h**) Number of calcium transients showing Δ*F/F*_0_ > 0.04 in 5 dpf Tg[Huc:GCaMP5G] larvae at different time points following exposure to either 15 µM DFP (*n* = 4) or vehicle (*n* = 3) (two-way ANOVA with Sidak’s multiple comparisons test: n.s., non-significant; **, *p* < 0.01; ****, *p* < 0.0001). (**i**) Pattern of calcium transients seen in 5 dpf Tg[Huc:GCaMP5G] larvae exposed for 5 h to 15 µM DFP and then to 15 µM DFP + 40 µM diazepam (DZP) for an additional hour. (**j**) Number of calcium transients showing Δ*F/F*_0_ > 0.04 in 5 dpf Tg[Huc:GCaMP5G] larvae exposed for 5 h to 15 µM DFP and then to 15 µM DFP + 40 µM diazepam (DZP) for an additional hour (*n* = 6) (Student’s unpaired *t*-test: **, *p* < 0.01).

### DFP exposure induces increased neuronal apoptosis

Exposure to warfare OP nerve agents, such as soman, sarin, or VX, causes massive neuronal loss in both humans and animal models^28–32^. We therefore examined whether DFP exposure also caused increased neuronal death, using both *in vivo* acridine orange (AO) staining, a vital marker that labels fragmented DNA in apoptotic cells, and immunodetection of activated-caspase-3 (Act-casp3), a marker of early apoptosis stages (Fig. **4a, b**). Interestingly, DFP-exposed larvae displayed a significantly increased number of neurons expressing Act-casp3 compared to that observed in controls (Fig. **4c**-**e**). In addition, *in vivo* AO labeling confirmed that DFP exposure caused an increased neuronal apoptosis (Fig. **3f**-**i**). We then investigated whether the increased number of apoptotic neurons seen in DFP-treated larvae was a direct consequence of neuronal hyperexcitation. To this end, 5 dpf larvae were co-treated for 6 h with 15 µM DFP and 40 µM DZP, and neuronal apoptosis was visualized by AO labeling. Results indicate that DZP was able to partially alleviate DFP-induced neuronal apoptosis (Fig. **3h, i**), strongly suggesting a link between neuronal hyperexcitation and neuronal death.

**Fig. 4.**
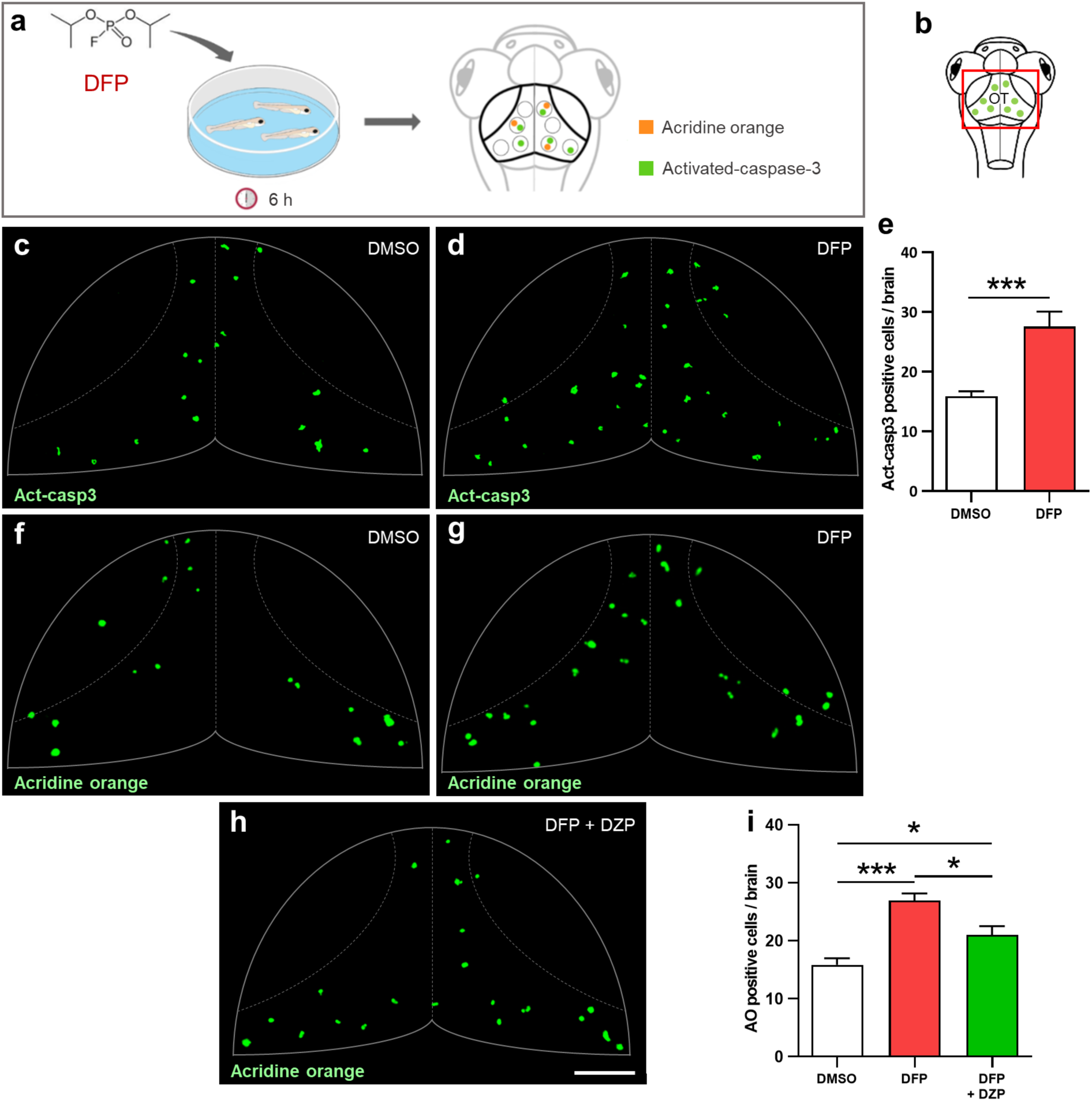
DFP exposure increased neuronal apoptosis. (**a)** As the experimental set-up, 5 dpf larvae were exposed to either 15 µM DFP or vehicle (1% DMSO) for 6 h, before processing for either acridine orange (AO) staining or anti-activated caspase-3 (Act-casp3) immunolabeling. (**b**) Scheme of a 5 dpf larva’s head with the red box showing the region of interest in the brain, uncovering the optic tectum (OT). (**c, d**) Act-casp3 immunolabeling of OT neurons in 5 dpf larvae exposed for 6 h to either vehicle (**c**) or 15 µM DFP (**d**). (**e**) Quantification of Act-casp3-positive neurons in 5 dpf larvae exposed for 6 h to either 15 µM DFP (*n* = 12) or vehicle (*n* = 12) (Student’s unpaired *t*-test with Welch’s correction: ***, *p* < 0.001). (**f-h**) Visualization of AO-labeled apoptotic neurons in 5 dpf larvae exposed for 6 h to either vehicle (**f**), or 15 µM DFP (**g**) or 15 µM DFP + 40 µM diazepam (**h**). (**i)** Quantification of the number of acridine orange positive cells in 5 dpf larvae exposed for 6 h to either vehicle (*n* = 24), or 15 µM DFP (*n* = 17) or 15 µM DFP + 40 µM diazepam (DZP) (*n* = 10) (one-way ANOVA with Tukey’s multiple comparisons test: *, *p* < 0.05; ***, *p* < 0.001). Scale bar: 50 µm.

### Increased NR2B-NMDA subunit receptor expression and decreased GAD65/67 and gephyrin protein accumulation in DFP exposed larvae

It has been shown that following OP exposure, neuronal seizures are associated with both an increased glutamatergic response and excessive NMDA receptor activation^33–35^. To examine whether accumulation of the NR2B-NMDA subunit receptor, a component of the main excitatory glutamate receptor, was modified following DFP exposure, brain sections of larvae exposed to DFP or controls were analyzed by immunocytochemistry using an anti-NR2B-NMDA antibody (Fig. **5a, b**). Interestingly, results showed a clear increase in the accumulation of NR2B-NMDA protein in the tectal neuropil of DFP-exposed larvae, compared to that seen in controls (Fig. **5 c, d**). Moreover, the DFP-induced increased NR2B-NMDA accumulation at excitatory synapses was confirmed by quantification of synaptic puncta in the tectal neuropil (Fig. **5e**). We also analyzed the synaptic protein gephyrin, which l is a scaffolding protein expressed at the inhibitory post-synaptic terminals (Fig. **5a, f**). Quantification of this labeling showed a marked decrease in gephyrin synaptic puncta density in the tectal neuropil in DFP-exposed larvae compared to controls (Fig. **5g**-**i**). To further characterize brain reorganization induced by DFP exposure, we analyzed, by immunohistochemistry, the accumulation of glutamate decarboxylase 65/67 (GAD65/67), the enzymes involved in GABA synthesis, the main inhibitory transmitter (Fig. **5a, j**). Interestingly, quantification of immunolabeling signals indicated that DFP-treated larvae displayed markedly decreased accumulation of GAD65/67 proteins (Fig. **5k-m**). Altogether, these results strongly suggest that DFP exposure induces neuronal hyperexcitation as a result of simultaneous increased glutamatergic activity and decreased GABAergic signaling.

**Fig. 5.**
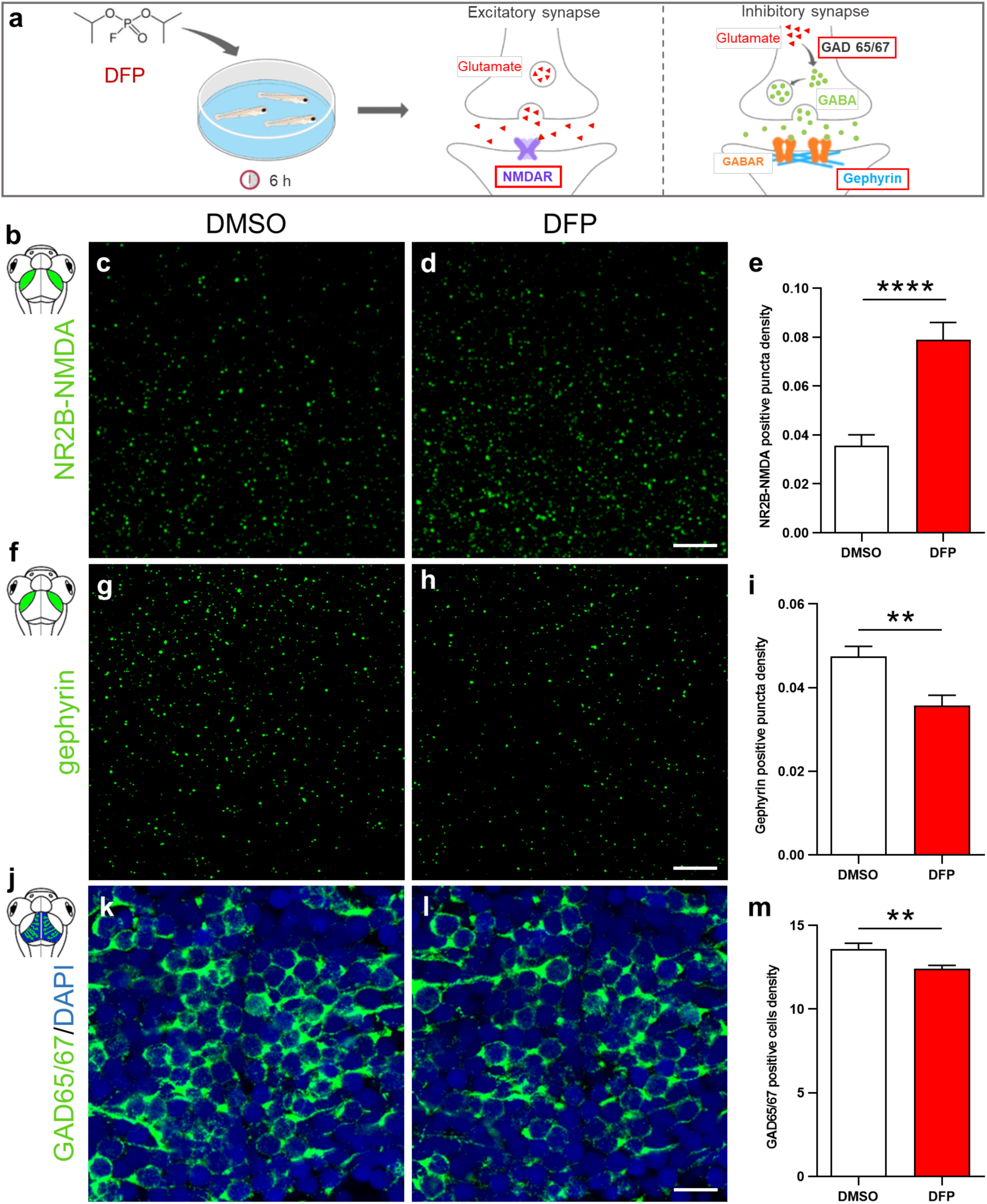
DFP exposure caused increased NR2B-NMDA subunit receptor accumulation combined with decreased gephyrin and GABA signaling. (**a**) As experimental set-up, 5 dpf larvae were exposed to either 15 µM DFP or vehicle (DMSO) for 6 hours, prior to being processed for NR2B-NMDA, gephyrin or GAD 65/67 immunolabeling. (**b**) Scheme of 5 dpf larval head highlighting the tectal neuropils in green. (**c-d**) NR2B-NMDA receptor immunolabelling of 5 dpf larvae brains exposed to either DMSO (**c**) or 15 µM DFP (**d**). Scale bar: 5 µm. (**e**) Quantification of NR2B-NMDA puncta density in 5 dpf larvae treated for 6 hours with either DMSO (*N* = 4; *n* = 22) or 15 µM DFP (*N* = 4; *n* = 26) (Mann Whitney : ****, *P* < 0.0001). (**f**) Scheme of 5 dpf larval head highlighting the tectal neuropils in green. (**g-h**) Gephyrin immunolabelling of 5 dpf larvae brains exposed to either DMSO (**g**) or 15 µM DFP (**h**). Scale bar: 5µm. (**i**) Quantification of gephyrin puncta density in 5 dpf larvae treated for 6 hours with either DMSO (*N* = 4; *n* = 18) or 15 µM DFP (*N* = 4; *n* = 18) (Student unpaired *t*-test: **, *P* < 0.01). (**j**) Scheme of 5 dpf larval head highlighting stratum periventriculare in blue and green. (**k-l**) GAD65/67 immunolabelling of neurons in the optic tectum of 5 dpf larvae brains exposed to either DMSO (**k**) or 15 µM DFP (**l**). Scale bar :10 µm. (**m**) Quantification of the density of neurons expressing GAD65/67 protein in the optic tectum of 5 dpf larvae exposed to either DMSO (*N* = 4; *n* = 18) or 15 µM DFP (*N* = 4; *n* = 19) (Student unpaired *t*-test : **, *P* < 0.01). *N* = number of larvae and *n* = number of slices.

## Discussion

Because of the widespread use of organophosphorus (OP) compounds for agricultural purposes and the lack of fully effective countermeasures, OP poisoning remains a major public health issue worldwide, with several million intoxications and over 200,000 deaths reported each year^36,37^, emphasizing the need for fully efficient antidotes to alleviate OP toxicity. To help fill this gap, we used the possibilities offered by zebrafish larvae and developed a vertebrate model of OP poisoning to investigate the consequences of OP exposure on neuronal functions and neuronal network activity. As an OP compound, we chose DFP, a moderately toxic compound previously used for OP toxicology research^38^. In good agreement with the results gained with rodent OP poisoning models^39,40^, zebrafish larvae exposed to DFP displayed marked AChE inhibition and muscle paralysis, two important hallmarks of OP poisoning. More importantly, unlike rodent OP intoxication models, in which premature death due to respiratory failure must imperatively be prevented by the simultaneous addition of low dose atropine and oximes, there is no need to treat DFP-exposed zebrafish with acetylcholine modulators, making DFP-exposed zebrafish larvae an accurate animal model to investigate the physiopathology of OP poisoning. In particular, as a pure OP poisoning model, zebrafish larvae also offer a powerful tool to test the effects of anti-convulsive agents or other OP antidotes, in the absence of either muscarinic antagonists or cholinesterase reactivators.

Previous results in both humans and rodent models have shown that acute OP poisoning causes neuronal hyperexcitation leading to epileptic seizures^40^. In good agreement with this observation, we first observed that zebrafish larvae exposed to DFP displayed a considerably increased accumulation of the Fosab protein in brain neurons. *c-Fos* is a member of the IEG family, widely used as a molecular marker of neuronal activity, and overexpressed in a large number of animal epilepsy models^41–43^, suggesting that acute DFP exposure causes neuronal hyperexcitation. Using qRT-PCR analysis of brain RNAs, we next showed that DFP exposure also induced a massive accumulation of transcripts of the IEGs *fosab, atf3, junBa*, and *npas4b*, providing another indication that DFP induces a massive neuronal hyperexcitation in zebrafish larvae. To further investigate whether DFP exposure causes neuronal hyperexcitation, we used calcium imaging, a recent technology that enables the visualization of neuronal excitation and epileptic seizures in living zebrafish larvae^14,26,44^. Interestingly, in larvae exposed to acute DFP poisoning, we observed a massive increase in the number of neuronal calcium transients compared to those seen in controls, and also a significant increase in the amplitude of the transients in DFP-treated larvae. Last, these calcium transients detected in brain neurons of DFP-exposed larvae were significantly diminished after addition of diazepam, a GABA receptor agonist. Altogether, these observations demonstrate that after acute DFP poisoning, zebrafish larvae display epileptiform neuronal hyperexcitation. In humans suffering from acute OP intoxication, whenever neuronal hyperexcitation is not rapidly treated, epileptic seizures eventually develop into SE, a major life-threatening neurological condition^6,45^, which also causes long-term brain damage^46,47^. Interestingly, in the zebrafish model of OP poisoning presented here, we observed that almost all the larvae displayed numerous massive calcium transients from 2 to 3 h after DFP exposure onward, which were strongly reminiscent of those associated with generalized seizures seen in both genetic and pharmacological zebrafish epilepsy models^14,26^. Altogether, these findings strongly suggest that DFP exposure causes epileptiform neuronal hyperexcitation in zebrafish larvae, eventually leading to a status epilepticus-like condition. This validated DFP model could be used to identify novel OP antidotes, and especially neuroprotective entities preventing the long-lasting neuronal morbidities associated with OP exposure. In particular, new AChE reactivators or other efficient countermeasures could be identified on the basis of their ability to restore locomotor activity or counteract epileptogenic effects of OP poisoning.

Together with AChE inhibition and epileptic seizures, massive neuronal death is another hallmark of OP poisoning^23^, also observed in DFP-exposed zebrafish larvae, as demonstrated by acridine orange staining and Act-casp3 immunocytochemistry. Importantly, given that the number of Act-casp3 positive cells was approximately doubled only 6 h after OP addition, and that the cleavage of Pro-Caspase 3 into Act-casp3 is a mid-stage event in the intrinsic apoptotic cascade, these data suggest that DFP-induced neuronal apoptosis is initiated in the very first hours that follow OP poisoning, a time frame also matching the massive calcium transients seen in DFP-treated individuals. Altogether, these results suggest that the increased rate of neuronal apoptosis seen in DFP-treated individuals was a consequence of glutamate excitotoxicity due to epileptiform neuronal hyperexcitation. Alterations in proteolytic cleavage of pro-caspase 1 and 3 have already been described in different experimental epilepsy models^48,49^. However, the relationship between neuronal excitation, glutamate excitotoxicity and neuronal apoptosis remains poorly understood. At the cellular level, it has long been known that epileptiform neuronal hyperexcitation relies on massive glutamate releases, inducing glutamate receptor over-activation at glutamatergic excitatory synapses^33–35^. In the case of OP poisoning, it has been shown that acute OP intoxication induces activation of NMDA receptors^50^, which in turn causes neuronal seizures and apoptosis^46,51^. Interestingly, in the model of OP poisoning presented here, hyperexcitation of brain neurons was correlated with both increased accumulation of the NR2B subunit of excitatory NMDA receptor and decreased accumulation of gephyrin and GAD65/67 proteins, two important components of GABAergic inhibitory signaling. These results suggest that after DFP poisoning, neuronal hyperexcitation is due to a shift in the glutamatergic/GABAergic activity balance of brain neurons toward excitatory states.

We report here on a zebrafish model of OP poisoning that faithfully recapitulates the neuronal deficits observed in humans, including AChE inhibition, epileptiform neuronal hyperexcitation, and neuronal apoptosis. Moreover, this vertebrate model does not require the simultaneous addition of acetylcholine modulators, unlike rodent models of OP poisoning, thus providing a pure and accurate model for large-scale *in vivo* screening of entities that could restore CNS functions after OP poisoning and alleviate the long-term neurological sequelae of such intoxication.

## Materials and methods

### Fish husbandry and zebrafish lines

Zebrafish were kept at 26–28 °C in a 14 h light/10 h dark cycle. Embryos were collected by natural spawning and raised in E3 solution at 28.5 °C. To inhibit embryo pigmentation, 0.003% 1-phenyl-2-thiourea was added at 1-day post-fertilization (dpf). Wild-type AB fish, originally purchased from Zebrafish International Resource Center (Eugene, OR, USA) or Tg[HuC:GCaMP5G] transgenic line^52^, was a gift from Dr. George Debrégeas (Laboratoire Jean Perrin, Paris). These lines were raised in our facility. All animal experiments were conducted at the French National Institute of Health and Medical Research (Inserm) UMR 1141 in Paris in accordance with European Union guidelines for the handling of laboratory animals (http://ec.europa.eu/environment/chemicals/lab_animals/home_en.htm), and were approved by the Direction Départementale de la Protection des Populations de Paris and the French Animal Ethics Committee under reference No. 2012-15/676-0069.

### DFP treatment

Diisopropylfluorophosphate (DFP) was purchased from Sigma Aldrich. A stock solution (5.46 mM), stored at −20 °C, was diluted extemporaneously to 15 µM in 1% DMSO/E3 medium. Control zebrafish larvae were treated with 1% DMSO / E3 medium.

### Measurement of DFP stability

(See supplementary Materials & Methods)

### Morphological analysis

4 dpf larvae were anesthetized and lateral views of the whole body were acquired using the same parameters for all larvae. Body length, eye size, and head size were then measured using the ImageJ software. Body length was measured from the anterior tip of the body to the caudal peduncle. Eye size and brain surface area were measured by specifying the eye and brain boundaries.

### Hematoxylin/eosin staining

(See supplementary Materials & Methods)

### Measurement of AChE activity

(See supplementary Materials & Methods)

### Zebrafish larval locomotor activity

Larvae locomotor activity was measured using a Zebrabox (View Point), an automated infrared tracking device, with ZebraLab 5.13.0.240 software (https://www.viewpoint.fr/en/home). 5 dpf larvae were individually dispatched in a 96-well plate in approximately 200 µl of E3 medium containing either 1% DMSO or 15 µM DFP. The plate was then placed in the recording chamber for a 30-minute habituation in the dark and in silence. Locomotor activity was then recorded for 4 h using the following settings: animal color was set to black and threshold detection to 12. The locomotion activity was quantified as the sum of all pixels showing intensity changes during the recording time and plotted as “acting units”.

### qRT-PCR

For brain RNA isolation, larvae brains were dissected, and homogenized using a syringe equipped with a 26G needle (10 brains per sample) using the RNA XS Plus kit (Qiagen, Hilden, Germany). cDNAs were synthesized using the iScript cDNA Synthesis Kit (Bio-Rad, Munich, Germany) and qPCR was performed using iQ SYBR Green Supermix (Bio-Rad). Samples were run in triplicate. Expression levels were normalized to that of the *tbp* gene. The primers (Eurofins Genomics, Ebersberg, Germany) used are listed in Supplementary Table 1.

### Neuronal calcium transient imaging

5 dpf larvae were paralyzed using 300 µM pancuronium bromide (PB) and embedded in 1.3% low-melting agarose in the center of a 35 mm glass-bottomed dish, and then covered with 3 mL of E3 solution containing 300 µM PB. The recording chamber was then placed under a Leica SP8 laser scanning confocal microscope equipped with a 20×/0.75 multi-immersion objective. Vehicle (1% DMSO) or DFP (15 µM) was added to the E3 medium and calcium activity was recorded for 6 h. Calcium fluorescent signals were recorded on a single focal plan, located approximately in the middle of the optic tectum, at a 512 × 512-pixel image resolution and a frame rate of 2 images per second. Fluorescence intensity of each frame was measured using a homemade macro on ImageJ 1.52p (https://imagej.nih.gov/ij/). Fluorescence variations (Δ*F/F*_0_) were calculated using Microsoft Excel (for Windows 2013, version 15.0.4569.1506) by subtracting the mean fluorescence intensity of all frames (*F*_0_) and dividing the results by *F*_0_. A subtraction of the mean value of the lowest (all values under the median) within a 20 s sliding window around the point was finally applied to correct the fluorescence drift occurring during long calcium recordings. Fluorescence variations greater than 0.04 Δ*F/F*_0_ were considered as calcium events upon visual confirmation since the detection system may detect false events.

### Diazepam treatment

5 dpf Tg[Huc:GCaMP5G] larvae were exposed to 15 µM DFP for 5 hours, PB-paralyzed and embedded in 1.1% low-melting agarose in the center of a 35 mm glass-bottomed dish, and then covered with an E3 solution containing 15 µM DFP and 300 µM PB. Calcium transients were first monitored for 30 min. Diazepam (40 µM DZP, Sigma) was then added and calcium transients were recorded for an additional hour. Visualization and recording of calcium transients were carried out as described above.

### Acridine orange labeling of apoptotic cells

To quantify neuronal cell death, we used *in vivo* acridine orange (AO) staining of fragmented DNA molecules in apoptotic cells. 5 dpf larvae treated with either vehicle or DFP were incubated for 30 minutes in AO (1/2000, VectaCell), thoroughly washed several times, PB-paralyzed and agar-embedded. 120 µm stacks of brain sections were then acquired using a Leica SP8 laser scanning confocal microscope equipped with a 20×/0.75 multi-immersion objective. In addition to AO staining, brains were stained with anti-activated-caspase-3 and processed as previously described (See supplementary Materials & Methods).

### Immunohistochemistry (see also Supplementary Table 2 for details on antibodies used)

For protein immunodetection, zebrafish larvae were fixed in 4% formaldehyde, incubated overnight in 15% sucrose at 4 °C, embedded in 7.5% gelatin/15% sucrose solution, flash frozen in isopentane at −45 °C and stored at −80 °C until use. For gephyrin immunodetection, unfixed larvae were directly incubated for 30 minutes in 15% sucrose at room temperature and then embedded in 7.5% gelatin/15% sucrose solution, flash frozen in isopentane at −45 °C and stored at −80 °C until use. Frozen embedded larvae were cut into 20 µm cryostat sections. For Fosab staining, we strictly applied the protocol described by Chatterjee et al. (2015)^53^. For anti-GAD56/67 and anti-gephyrin immunolabeling, we used the endogenous biotin blocking procedure (Avidin/Biotin Blocking Kit, Dako, code No. X0590) according to the manufacturer’s instructions. Sections were then blocked and permeabilized with 0.2% gelatine, 0.25% Triton X100 diluted in 1X PBS for 1 h at room temperature and then incubated overnight at room temperature with either anti-GAD65/67 (1:300) or anti-gephyrin (1:100). After several washes, GAD65/67 and gephyrin proteins were detected using biotinylated goat anti-rabbit and streptavidine Alexa 488 (Molecular Probes; catalog No. S32355; used at 1:400 dilution). Sections were counterstained for 10 minutes with 0.1% DAPI (Sigma, St. Louis, MO). For anti-NMDA-NR2B, immunohistochemistry was performed as previously described^54^. Briefly, brain tissue sections were blocked and permeabilized with 0.2% gelatin, 0.25% Triton X100 diluted in PBS for 1 h at room temperature and then incubated overnight at room temperature with anti-NMDA-NR2B antibody (1:300). After several washes, sections were incubated for 1 h with anti-rabbit coupled to Alexa Fluor 488. Sections were counterstained for 10 minutes with DAPI (Sigma-Aldrich, used at 1:3000) before mounting.

### Quantification of NR2B and gephyrin puncta labeling

Sections stained with anti-NR2B-NMDA and anti-gephyrin antibodies were imaged at full resolution (voxel size: 0.063 × 0.063 × 0.3 µm) using a Leica SP8 laser scanning confocal microscope equipped with a 63x/1.4 oil-immersion objective. Images were then processed with AutoQuant X3.1 software (http://www.mediacy.com/autoquantx3) and the density of NR2B or gephyrin labeled puncta was quantified using a homemade ImageJ macro (Zsolt Csaba, Inserm UMR1141). After stacking all images, a median filter was applied, a threshold on the fluorescence intensity was then set to select NR2B-NMDA or gephyrin puncta. Puncta were identified as accumulation of more than two pixels (0.126 µm) in diameter, which lead to an average punctum size of 0.34 µm. The density of NR2B-NMDA or gephyrin puncta were calculated by dividing the number of labeled puncta detected in the tectal neuropil by the surface area of this region.

### Quantification of Fosab- and GAD65/67-positive neuron density

Sections hybridized with anti-Fosab and anti-GAD65/67 antibodies were imaged using a Leica SP8 laser scanning confocal microscope equipped with a 40×/1.3 oil-immersion objective. The neurons expressing Fosab or GAD65/67 were manually counted on sections of optic tectum or telencephalon. The number of positive neurons was then divided by the surface area of the corresponding regions to calculate a density of neurons expressing those proteins in larvae exposed to DFP or vehicle.

### Statistical analysis

Statistical analyses were performed using GraphPad Prism 8.4.3.686 (https://www.graphpad.com/scientific-software/prism/). Data were first challenged for normality using the Shapiro-Wilk test. Data with a normal distribution were analyzed by a two-tailed unpaired *t*-test. Data not showing normal distribution were analyzed using a two-tailed Mann-Whitney test. All graphs show mean ± s.e.m.

## Supporting information

Supplementary Video 1. 3 minute-long representative recording of calcium activity imaging in optic tectum neurons of 5 dpf larvae following 3 h exposu

Supplementary Video 2. 3 minute-long representative recording of calcium activity imaging in optic tectum neurons of 5 dpf larvae following 3 h exposi

## Data availability

The datasets generated during the current study are available from the corresponding author on reasonable request.

## Author contributions

A.B. and J.S. performed the experiments, the analysis, and designed the figures. R.H.A., A.I., D.S., C.R., O.B. and N.T. helped carry out the experiments. N.D. helped supervise the project. C.Y., G.D.B., F.N. and N.D. contributed to the final manuscript. N.S.Y. supervised the project and wrote the manuscript.

## Competing interests

The authors declare no competing interest.

## Notes

### Competing Interest Statement

The authors have declared no competing interest.

### Summary of Updates

With millions of intoxications each year and over 200,000 deaths, organophosphorus (OP) compounds are an important public health issue worldwide. OP poisoning induces cholinergic syndrome, with muscle weakness, hypertension, and neuron damage that may lead to epileptic seizures and permanent psychomotor deficits. Existing countermeasures are lifesaving but do not prevent long-lasting neuronal comorbidities, emphasizing the urgent need for animal models to better understand OP neurotoxicity and identify novel antidotes. Here, using diisopropylfluorophosphate (DFP), a prototypic and moderately toxic OP, combined with zebrafish larvae, we first showed that DFP poisoning caused major acetylcholinesterase inhibition, resulting in paralysis and CNS neuron hyperactivation, as indicated by increased neuronal calcium transients and overexpression of the immediate early genes fosab, junBa, npas4b, and atf3. In addition to these epileptiform seizure-like events, DFP-exposed larvae showed increased neuronal apoptosis, which were both partially alleviated by diazepam treatment, suggesting a causal link between neuronal hyperexcitation and cell death. Last, DFP poisoning induced an altered balance of glutamatergic/GABAergic synaptic activity with increased NR2B-NMDA receptor accumulation combined with decreased GAD65/67 and gephyrin protein accumulation. The zebrafish DFP model presented here thus provides important novel insights into the pathophysiology of OP intoxication, making it a promising model to identify novel antidotes.

